# Deep proteome profiling of white adipose tissue reveals marked conservation and distinct features between different anatomical depots

**DOI:** 10.1101/2022.08.23.504892

**Authors:** Søren Madsen, Marin E. Nelson, Vinita Deshpande, Sean J. Humphrey, Kristen C. Cooke, Anna Howell, Alexis Diaz-Vegas, James G. Burchfield, Jacqueline Stöckli, David E. James

## Abstract

White adipose tissue is deposited mainly as subcutaneous adipose tissue (SAT), often associated with metabolic protection, and abdominal/visceral adipose tissue (VAT), which contributes to metabolic disease. To investigate the molecular underpinnings of these differences, we conducted comprehensive proteomics profiling of whole tissue and isolated adipocytes from these two depots across two diets from C57Bl/6J mice. The adipocyte proteomes from lean mice were highly conserved between depots, with the major depot-specific differences encoded by just 3% of the proteome. Adipocytes from SAT (SAdi) were enriched in pathways related to mitochondrial complex I and beiging, whereas visceral adipocytes (VAdi) were enriched in structural proteins and positive regulators of mTOR presumably to promote nutrient storage and cellular expansion. This indicates that SAdi are geared toward higher catabolic activity, while VAdi are more suited for lipid storage.

By comparing adipocytes from mice fed chow or Western diet (WD), we define a core adaptive proteomics signature consisting of increased extracellular matrix proteins and decreased fatty acid metabolism and mitochondrial Coenzyme Q biosynthesis. Relative to SAdi, VAdi displayed greater changes with WD including a pronounced decrease in mitochondrial proteins concomitant with upregulation of apoptotic signaling and decreased mitophagy, indicating pervasive mitochondrial stress. Furthermore, WD caused reduction in lipid handling and glucose uptake pathways particularly in VAdi, consistent with adipocyte de-differentiation. By overlaying the proteomics changes with diet in whole adipose tissue and isolated adipocytes, we uncovered concordance between adipocytes and tissue only in the VAT, indicating a unique tissue-specific adaptation to sustained WD in SAT.

Finally, an in-depth comparison of isolated adipocytes and 3T3-L1 proteomes revealed a high degree of overlap, supporting the utility of the 3T3-L1 adipocyte model. These deep proteomes provide an invaluable resource highlighting differences between white adipose depots that may fine-tune their unique functions and adaptation to an obesogenic environment.

## INTRODUCTION

White adipose tissue is one of the most adaptive tissues in mammals and can expand to account for greater than 70% of total body mass in extreme cases of sustained positive energy balance. Adipose tissue expandability is crucial to accommodate the storage of excess lipids in times of plenty and mobilize nutrients for use by tissues throughout the body in times of limited food availability. However, in the case of sustained positive energy balance, adipose tissue stores can be overwhelmed, resulting in spill over and accumulation of harmful ectopic lipids in other tissues such as cardiovascular tissue, skeletal muscle, liver and the pancreas. Intriguingly, in humans there is a strong relationship between visceral adiposity and metabolic disease risk (Fox et al., 2007). Conversely, individuals with a preponderance of subcutaneous fat are often protected from metabolic disease. Many theories have been proposed to explain these observations. Subcutaneous fat has higher neural innervation and, as a consequence, is enriched with “beige” adipocytes, which have elevated thermogenic capacity and so are protective against excess weight gain (Ahn et al., 2019; Contreras et al., 2014; Kajimura et al., 2015). Visceral fat, on the other hand, has higher infiltration of immune cells such as macrophages particularly in response to obesogenic environments (Ahn et al., 2019; Poret et al., 2018), which is associated with systemic inflammation and insulin resistance in other tissues (Zatterale et al., 2019). Therefore, it will be critical to understand the molecular underpinnings that confer depot specificity.

Gene expression analysis has yielded important insights into key differences between adipose depots (Jones et al., 2020; Soccio et al., 2017); however, mRNA and protein levels display considerable discordance (Vogel and Marcotte, 2012). Thus, it is critical to investigate adipose tissue composition at the proteome level to uncover depot-specific biology. It is unclear which aspects of the proteome define adipocytes from different depots in a lean, healthy context, or how different adipocyte proteomes adapt to obesogenic conditions. One possibility is that the proteomes from adipocytes from different depots are highly conserved and functional differences may be conferred by interactions with the microenvironment established by the surrounding stromal vascular cells. Since adipose tissue is a milieu of many cell types including fibroblasts and immune cells, it is crucial to compare both the adipocyte and whole tissue proteomes to link molecular differences between depots to physiologic consequences. Here we report a deep proteomics analysis of different depots using both whole tissue and isolated adipocytes from lean and diet-induced obese mice. In lean mice, we discovered that only 3% of the adipocyte proteome differs between the two depots, and we revealed that SAdi had enhanced capacity for catabolic processes, while VAdi were equipped for lipid storage and cell expansion. WD caused a greater divergence in the adipocyte proteomes, with the greatest changes occurring in visceral adipocytes and tissue compared to subcutaneous, including immune cell infiltration, downregulation of ‘adipocyte’ processes such as glucose metabolism, and upregulation of ‘fibroblastic’ processes including collagen deposition. Furthermore, we uncovered a pro-apoptotic proteomics signal in the VAdi after WD feeding pointing to severe mitochondrial dysfunction in these adipocytes. These data provide an invaluable adipose-centric resource for the metabolic research community by highlighting both similarities and key differences that emerge between the biology of each adipose depot, and how each depot adapts to overnutrition.

## Experimental Procedures

### Animals

C57BL/6J male mice were obtained from Australian BioResources (Moss Vale, NSW, Australia). Mice were maintained at 23°C on a 12-hour light-dark cycle and *ad libitum* access to food and water. From weaning, mice were fed standard laboratory chow (containing 12% calories from fat, 65% calories from carbohydrate, 23% calories from protein (‘Irradiated Rat and Mouse Diet’, Specialty Feeds, Glen Forest, WA, Australia)). For WD studies, mice were fed a diet made in-house containing 45% fat, 35% carbohydrate and 20% protein as we have described (Nelson et al., 2022) for 9 months from 11 to 14 weeks of age. Mice were weighed once per week. Body composition was analyzed at 43 weeks of age using quantitative magnetic resonance technology (EchoMRI Body Composition Analyser, EchoMRI). For glucose tolerance tests, 43-week-old mice were fasted for 6 hours and received an oral bolus of glucose (1 g/kg lean mass). Blood was sampled from the tail vein at indicated time points using an Accu-Check II glucometer (Roche Diagnostics). For insulin measurements, whole blood was collected from the tail vein at basal and 15 minutes after oral glucose into wells of commercially available ELISA kit (Crystal Chem, IL, USA) containing sample buffer, then the assay was carried out according to the manufacturer’s specifications. Results were multiplied by a factor of 2 to estimate the concentration of insulin in the plasma. For histological assessment of white adipose tissue, 8-10 week old C57BL/6J mice were fed either chow or WD for eight weeks. Experiments were performed in accordance with NHMRC (Australia) guidelines and under the approval of The University of Sydney Animal Ethics Committee.

### Tissue collection and primary adipocyte isolation

Mice were sacrificed by cervical dislocation. The epididymal fat pad was taken as ‘visceral’ adipose, and was excised carefully to avoid the testes. The inguinal fat pad was taken as ‘subcutaneous’ adipose, from which lymph nodes were removed following excision. Fat pads were rinsed in PBS and either snap frozen in liquid nitrogen or stored in fresh HEPES buffer (120 mM NaCl, 30 mM HEPES, 10 mM NaHCO_3_, 5 mM glucose, 4.7 mM KCl, 2 mM CaCl_2_, 1.18 mM KH_2_PO_4_, 1.17 mM MgSO_4_.7H_2_O, 1% BSA, pH 7.4) for immediate adipocyte isolation. The two diet groups were subdivided into three groups each and pooled for the isolation of primary adipocytes. Each group of mice (chow 1 (n=4), chow 2 (n=4), chow 3 (n=3); WD 1 (n=5), WD 2 (n=5), WD 3 (n=5)) represents one pooled data point for proteomics analysis. Adipose tissues were minced in HEPES buffer until pieces were approximately < 1 mm^2^ in size. Collagenase (Type I, Worthington) was added at 0.5 mg/mL for visceral and 1 mg/mL for subcutaneous adipose tissue, and digested for 1 hour at 37°C. Samples were passed through a 250 μM (chow adipocytes) or 300 μM (WD adipocytes) nylon mesh (Spectrum Labs) and washed 3 times with HES buffer (250 mM sucrose, 20 mM HEPES, 1 mM EDTA, pH 7.4). Between washes, adipocytes were left to float. Lysis buffer (2% sodium deoxycholate (Sigma), 200 mM Tris HCl, pH 8.5) was added and stored at −80°C until further processing.

### Cell culture – 3T3-L1 adipocytes

3T3-L1 fibroblasts (a gift from Howard Green, Harvard Medical School) were grown in Dulbecco’s modified Eagle’s medium (DMEM) containing 10% fetal bovine serum (Sigma) and 2 mM GlutaMAX (Gibco) in 10% CO_2_ at 37°C. For differentiation into adipocytes, cells were re-seeded and rapidly grown to confluence within 24h, then treated with DMEM/FBS containing 4 μg/ml insulin, 0.25 mM dexamethasone, 0.5 mM 3-isobutyl-1-methylxanthine and 100 ng/ml d-biotin. After 72 h, the differentiation medium was replaced with fresh FCS/DMEM containing 4 μg/ml insulin and 100 ng/ml d-biotin for a further 3 days, then replaced with fresh FCS/DMEM. Adipocytes were re-fed with FCS/DMEM every 48h and used for experiments 10 days after initiation of differentiation. Cells were routinely tested for mycoplasma. Prior to harvesting, 3T3-L1 adipocyte cell monolayers were washed 5x with ice-cold PBS.

### Sample preparation – 3T3-L1 adipocytes, adipocyte, and adipose tissue proteomes

Adipocytes were processed according to the in-StageTip protocol (Kulak et al., 2014). Briefly, samples were lysed in an equal volume of SDC lysis buffer (2% sodium deoxycholate (Sigma), 200 mM Tris HCl, pH 8.5) with boiling at 95°C for 5 mins with mixing at 1,000 rpm on a ThermoMixer (Eppendorf), cooled on ice, sonicated using a tip probe sonicator (1x 20 seconds, 90% output), then spun at 21,000g for 15 mins at 4°C. For tissue samples, 100 – 600 mg tissue was added to 1.5 mL lysis buffer, lysed using a tip probe sonicator (4-5x 20 seconds, 90% output), and spun at 21,000g for 30 mins at 4°C. After centrifugation, fat layers were carefully removed and discarded, and an aliquot was taken from which protein quantification performed using the Pierce BCA Protein Assay (Thermo Fisher Scientific). 60 μg of protein was extracted into clean tubes and samples topped with lysis buffer to obtain equal volumes. Proteins were reduced and alkylated with the addition of TCEP (Thermo Fisher Scientific) and CAA (Sigma) to final concentrations of 10 mM and 40 mM respectively at 95°C for 5 mins at 1,000 rpm on a ThermoMixer. Trypsin (Sigma) and LysC (Wako, Japan) were added in ratio of 1 μg enzyme to 50 μg protein, and samples digested at 37 °C overnight for 18 hours with mixing at 2,000 rpm on a ThermoMixer. Digested peptides were diluted 1:1 with water, and then desalted on SDBRPS StageTips as follows. Samples were diluted 50% with 99% EA (ethyl acetate)/1% TFA (trifluoracetic acid), vortexed thoroughly, and loaded onto StageTips packed with 2x disks SDBRPS material (3M Empore). StageTips were washed 1x with 100 μL 99% ethyl acetate/1% TFA, and 2x with 100 μL 0.2% TFA, then eluted with 5% ammonia/80% ACN (acetonitrile), and dried in a vacuum concentrator (Eppendorf) prior to fractionation.

### Offline peptide fractionation by StageTip-based SCX (adipocyte and adipose tissue samples)

Peptides derived from mouse adipocyte and adipose tissue samples were separated into 3 fractions using StageTip-based SCX fractionation (Ishihama et al., 2006). Briefly, approximately 30 μg of peptides were resuspended in 1% TFA, and loaded onto StageTips packed with 6x disks of SCX material (3M Empore). Peptides were eluted and collected separately with increasing concentrations of ammonium acetate (150 mM and 300 mM) in 20% ACN, followed by 5% ammonia/80% ACN. Collected peptide fractions were dried in a vacuum concentrator (Eppendorf) and resuspended in MS loading buffer (2% ACN/0.3% TFA).

### Offline peptide fractionation by high pH reversed-phase HPLC (3T3-L1 adipocytes)

Peptides derived from 3T3-L1 adipocytes were separated into 24 fractions using concatenated high pH reverse phase fractionation, as previously described (Yau et al., 2021). Briefly, peptides were fractionated using an UltiMate 3000 HPLC (Dionex, Thermo) with a XBridge Peptide BEH C18 column, (130A°, 3.5 mm 2.1 3 250 mm, Waters). 30 μg of peptides were resuspended in buffer A and loaded onto the column that was maintained at 50°C using a column oven. Buffer A comprised 10mM ammonium formate/2% ACN and buffer B 10 mM ammonium formate/80% ACN, and buffers were adjusted to pH 9.0 with ammonium hydroxide. Peptides were separated by a gradient of 10 - 40% buffer B over 4.4 min, followed by 40 - 100% buffer B over 1 min. Peptides were collected for a total duration of 6.4 min, with 72 fractions concatenated directly into 24 wells of a 96-well deep-well plate (3 concatenated fractions per well) using an automated fraction collector (Dionex, Thermo) maintained at 4°C. After fraction, samples were dried down directly in the deep-well plate and resuspended in MS loading buffer (2% ACN/0.3% TFA) prior to LC-MS analysis.

### Mass spectrometry analysis – adipocyte and adipose tissue proteomes

For the mouse adipocyte and adipose tissue proteomes, peptides were analyzed by mass spectrometry using a Dionex Ultimate 3000 UHPLC coupled to a Q Exactive Plus benchtop Orbitrap Mass Spectrometer (Thermo Fisher Scientific). Peptides were loaded onto an inhouse packed 75 μm ID x 50 cm column packed with 1.9 μm C18 material (Dr Maisch, ReproSil Pur C18-AQ), and separated with a gradient of 5-30% ACN containing 0.1% FA over 95 min at 300 nl/min, and column temperature was maintained at 60°C with a column oven (Sonation). MS1 scans were acquired from 300-1,650 m/z (35,000 resolution, 3e6 fill target, 20 ms maximum fill time), followed by MS/MS data-dependent acquisition of the top 15 ions using high-energy dissociation (HCD), with MS2 fragment ions read out in the Orbitrap (17,500 resolution, 1e5 AGC, 25 ms maximum fill time, 25 NCE, 1.4 m/z isolation width).

### Mass spectrometry analysis – 3T3-L1 adipocyte proteomes

For the 3T3-L1 proteome samples, peptides were analyzed by mass spectrometry using a Dionex Ultimate 3000 coupled to a Q Exactive HF-X mass spectrometer (Thermo Fisher Scientific). Peptides were loaded onto an in-house packed 75 μm ID x 50 cm column packed with 1.9 μm C18 material (DrMaisch, ReproSil Pur C18-AQ) maintained at 60°C with a column oven (Sonation). Peptides were eluted with a gradient of 5-30% Buffer B (80% ACN/0.1% Formic acid) over 40 min at a flow rate of 300 nL/min, and analyzed by data-dependent acquisition with one full scan (350 – 1,400 m/z; R = 60,000 at 200 m/z) at a target of 3e6 ions, followed by up to 20 data-dependent MS2 scans using HCD (target 1e5 ions; max. IT 28 ms; isolation window 1.4 m/z; NCE 27%; min. AGC target 1e4), detected in the Orbitrap mass analyser (R = 15,000 at 200 m/z). Dynamic exclusion (15 s) was switched on.

### Data Analysis

RAW MS data were processed using MaxQuant (version 1.5.9.1) (Cox and Mann, 2008) searched against a UniProt mouse database (January 2016 release). Default settings were used, and match between runs was switched on to facilitate the transfer of MS/MS identifications between equivalent and adjacent fraction measurements. The data was filtered to remove contaminants and proteins not quantified, and protein intensities were transformed to log2 scale.

### Data processing

Raw data were processed using MaxQuant (version 1.5.2.10) (Cox and Mann, 2008) searched against a UniProt mouse database (January 2016 release) using default settings with match between runs was switched on to facilitate the transfer of MS/MS identifications between equivalent and adjacent fraction measurements. Intensity values were normalized using total reporter area sum normalization. The data was then filtered to remove contaminants and proteins that were not quantified in any sample. To correct for a slight batch effect between adipose tissue and adipocytes, the tissue protein intensities were median normalized (**Fig. S1A**).

### Network analysis

To find associations between lipolysis, lipids synthesis and glucose uptake, STRING: Pubmed query in Cytoscape (v3.8.2) was used to identify the top 50 proteins within each term with a confidence score of 0.7 or greater. These three networks were then merged and filtered for differentially-regulated protein for both VAdi and SAdi.

### Estimating cell type proportions in whole adipose tissue proteomics

The R package BisqueRNA (Jew et al., 2020) was used for reference-based decomposition of the whole tissue proteomics data with ‘marker = NULL’ and ‘use.overlap = FALSE’. As reference, we utilized murine single RNA sequencing data (Emont et al., 2022). Seurat formatted data was downloaded (https://gitlab.com/rosen-lab/white-adipose-atlas) and data from all male mice was selected as input for cell type estimation.

### Gene set enrichment analysis

Gene set enrichment was performed with the web-based GEne SeT AnaLysis Toolkit (Liao et al., 2019) with the minimum number of genes in a pathway specified as 15 and maximal as 200 within the Gene Ontology Resource, or using the Reactome database (reactome.org). Pathways with p < 0.05 false discovery rate (FDR) were considered to be significantly overrepresented

### Experimental Design and Statistical Rationale

The aim of this study was to examine adipose depot adaptations to a sustained obesogenic challenge. To this end, the duration of feeding was selected based on prior time course data in C57Bl6/J fed similar diets (Burchfield et al., 2018). Adiposity reached a plateau at approximately 40 weeks (10 months) of WD feeding without changes to lean mass, indicating the time at which adipose storage capacity is exceeded. Therefore, we selected 9 months of WD feeding for this study to maximize WD expansion but before complete exhaustion of adipose fat storage capacity. To achieve measurement of the adipocyte proteome, the biological replicates analyzed comprised pooled isolated adipocytes from 3-5 mice. Pooling was necessary to ensure sufficient protein yields for proteomics analysis. “Enrichment” was considered at a fold change of ±2 and p < 0.05 unless otherwise stated. For histology, each group contained five animals, except for visceral adipose tissue from chow fed animals, which contained three animals.

### General data analysis

Bioinformatics, statistical analyses and plot generation were performed within the R statistical environment. Differential expression analysis was performed using the LIMMA package (Ritchie et al., 2015). Two-way ANOVAs were performed using a standard linear model function. All p values were adjusted for multiple testing using the Benjamini & Hochberg or Tukeys HSD method.

### Adipose tissue histology and adipocyte area

Formalin-fixed epididymal adipose tissue was paraffin embedded, sectioned, mounted on coverslips and stained with H&E. Coverslips were scanned to digital images using an Aperio ScanScope. Adipocyte cell area was then analyzed in ImageJ version 1.51 as described in (Nelson et al., 2022) with the following modifications. Images were converted to 8 bit and the threshold was adjusted so cell membranes were identified as signal and cell contents were identified as background in ImageJ version 1.51. Identification of cell membranes was performed in Ilastik (version 1.3.3) using supervised machine learning-based image segmentation. A sample of the images to be analyzed were selected as the training set and the object classification workflow was used to label regions of the images as object types (cell membrane, cytoplasm, or artifact). Using the uncertainty overlay, areas of high uncertainty were defined until the prediction layer showed satisfactory identification of the object types. The trained classifier was then run on all images and the object classification data was saved as simple segmentations of the original image in black (cytoplasm) and white (cell membranes). In ImageJ, the “Analyze Particles” built-in function was then applied to calculate cell areas with a defined size range of 400-80000 μm^2^ and a circularity range of 0.40-1.00. An entire cross section for each mouse was analyzed so that >5000 adipocytes were quantified per mouse.

## RESULTS

### Comprehensive deep proteome analysis of depot specific adipose tissue and adipocytes

We employed a deep proteomic analysis of whole-tissue and isolated adipocytes from subcutaneous and visceral white adipose depots of middle-aged C57Bl/6J mice fed chow or WD for 9 months. WD-fed mice (n=11) were obese and glucose intolerant compared to chow fed animals (n=15) (**Fig. S1B-F**). The cohort was divided into three biological replicates from each diet matched by metabolic parameters, allowing comparison of adipose tissue and adipocyte proteomes from two white adipose depots across two diets (**Fig. 1A**). A total of 7,359 proteins were quantified in at least one sample and 3,970 were quantified across all 24 samples (**Fig. 1B**). We quantified 6,372 and 5,051 proteins across all adipocyte or whole tissue proteomes, respectively. This difference can partly be explained by a greater missingness in whole tissue proteomes, where missing values ranged from 16-28% compared to 10-14% in the adipocyte proteomes (**Fig. S1G**). The overlap between the adipocyte and tissue proteomes was 4,789 proteins (**Fig. 1B**). Protein abundances spanned six orders of magnitude (**Fig. 1C**), making this proteome coverage the deepest reported for adipose tissue or adipocytes. Biological replicates were tightly correlated, with correlation coefficients above 0.93 (**Fig. S1H**). Principal component analyses revealed clustering of replicates, which separated by sample type (whole tissue and adipocytes) in principal component (PC) 1 and by diet in PC2 (**Fig. 1D**), highlighting a distinction between the whole tissue and adipocyte proteomes. Adipocyte markers and lipid droplet proteins, such as PLIN1, Perilipin-4 (PLIN4), Hormone Sensitive Lipase (HSL), Adipose triglyceride lipase (ATGL), and the insulin-dependent glucose transporter GLUT4 were all at least > 2 fold increased in the adipocyte proteome across both depots (**Fig 1E**), showing expected enrichment of parenchymal cells in the adipocyte proteome. Overall, ~1500 proteins showed > 2 fold increase in the adipocyte proteomes. These proteins were enriched for adipocyte-specific functions such as lipid and amino acid metabolism, as well as mitochondrial processes. Furthermore, PLIN1, Fatty Acid-Binding Protein (FABP4), Adiponectin (ADIPOQ) and Hormone Sensitive Lipase (HSL) were ranked in the top 3% most abundant adipocyte proteins.

**Figure 1.**
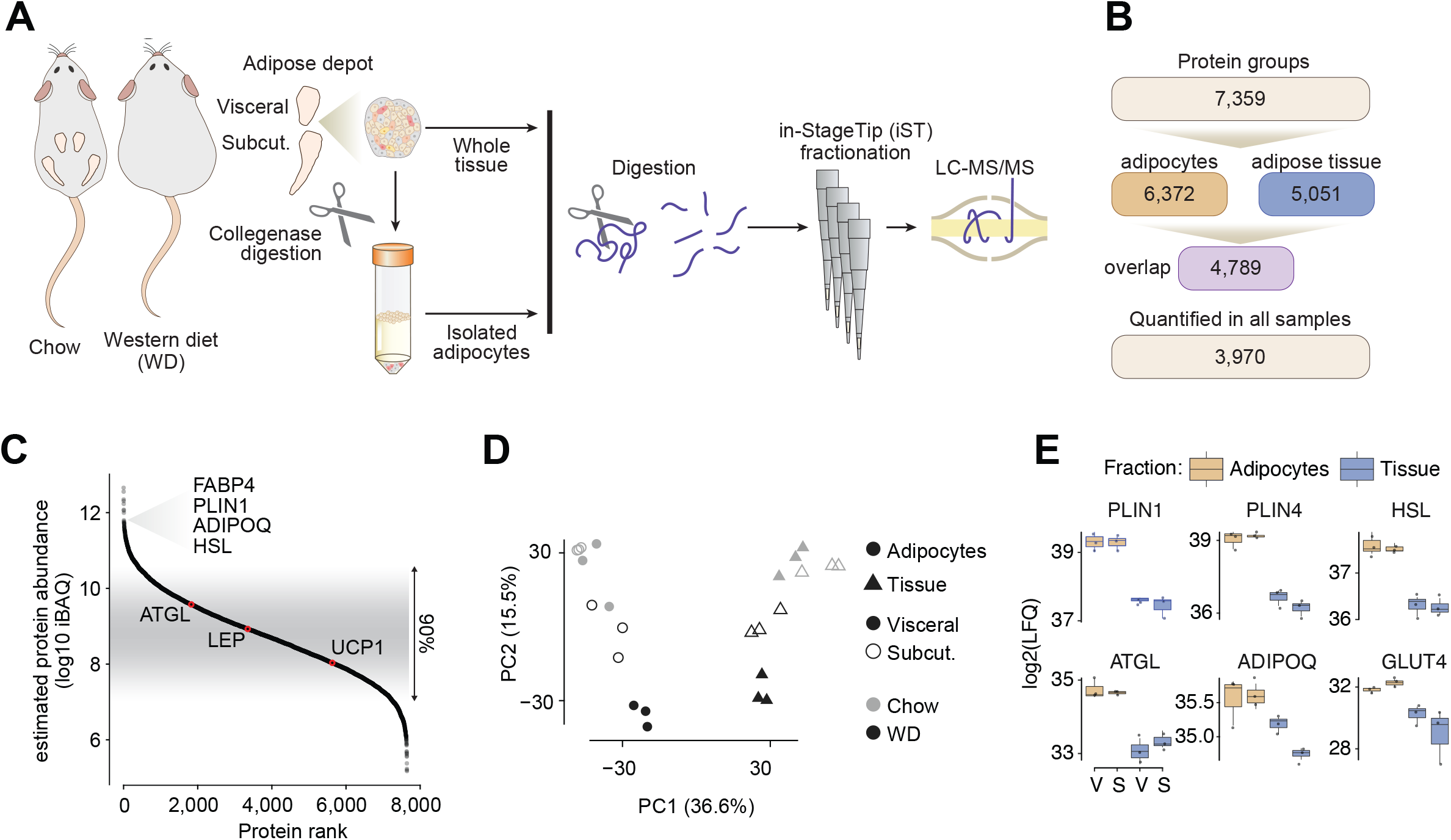
Deep proteomics coverage of adipose tissue and isolated adipocytes. **(A)** Schematic workflow for adipose tissue processing and proteomic analysis. **(B)** Quantification of proteins across proteomes. **(C)** Dynamic range of quantified proteins across all samples, based on intensity based absolute quantification (iBAQ). **(D)** Principal component analysis of across all samples. **(E)** Protein abundance of adipocyte markers in whole-tissue and isolated adipocytes from two visceral and subcutaneous adipose tissue from chow fed animals. V: visceral, S: subcutaneous.

### Adipocytes from different depots are strikingly similar under chow conditions

In chow-fed mice, we observed striking concordance between subcutaneous and visceral adipocytes (SAdi and VAdi, respectively) proteomes (R = 0.98; **Fig. 2A**). Remarkably, just 3% of the total proteome was different between adipocytes from the two depots, such that 146 proteins were exclusively identified in adipocytes from one depot and an additional 89 proteins were differentially expressed (**Fig. 2B**). Gene set enrichment analysis revealed that the most defining depot-specific features were an enrichment of select mitochondrial proteins in SAdi, particularly Complex I of the electron transport chain (ECT) and cytochrome complex assembly proteins, and an enrichment of cytoskeletal proteins in VAdi (**Fig. 2C**). These processes likely reflect differences in cell morphology and metabolic demand between the two depots.

**Figure 2.**
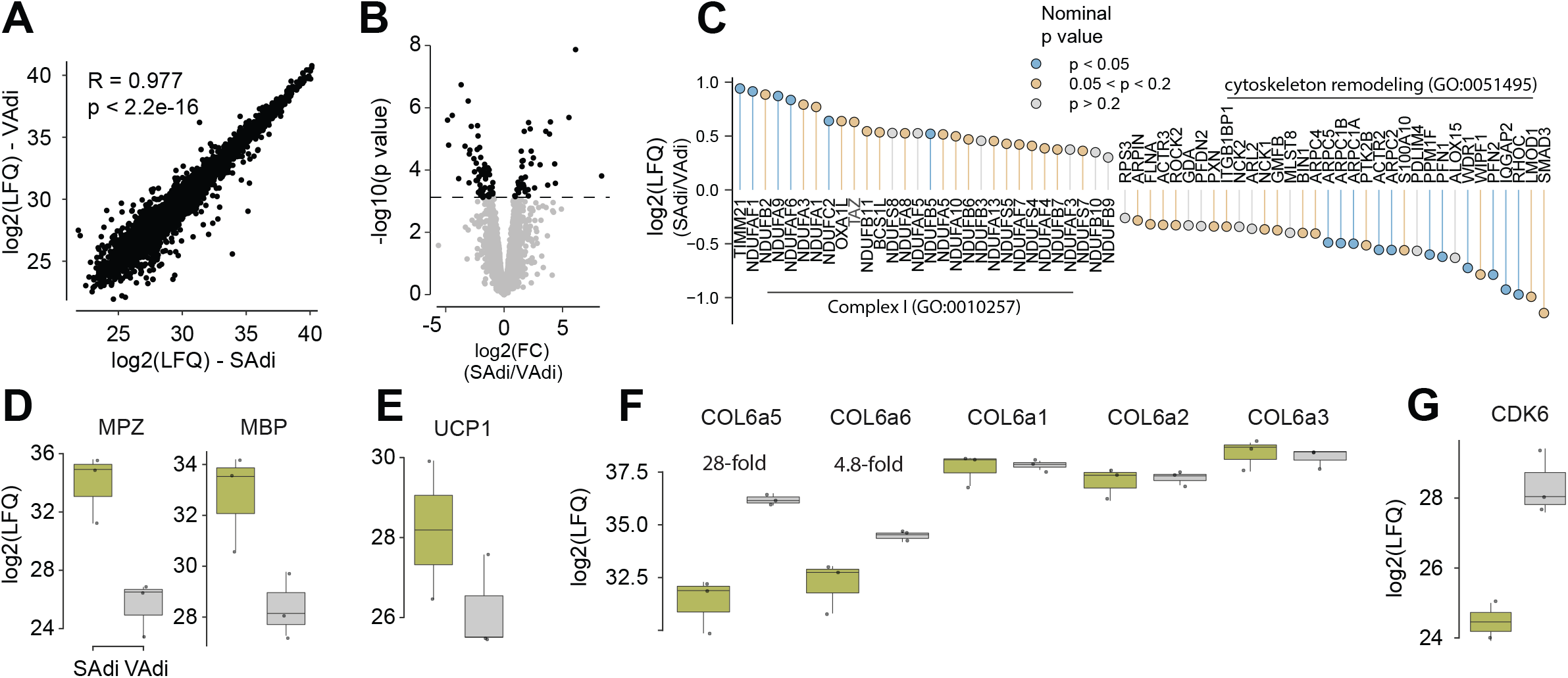
Comparison of isolated adipocytes from subcutaneous and visceral depots from chow fed animals. **(A)** Label-free quantitation (LFQ) of visceral adipocyte (VAdi) versus subcutaneous adipocyte (SAdi) proteomes. **(B)** Differences in protein abundances between VAdi and SAdi. **(C)** Fold-changes of proteins associated with mitochondrial Complex I and cytoskeletal remodeling. **(D-G)** Protein abundances of selected proteins. C: Chow, W: Western diet.

We identified 77 proteins exclusively expressed in SAdi, and only an additional 38 proteins that were significantly enriched compared to VAdi. Two proteins involved in Calcitonin-like ligand receptors signaling, Receptor Activity-Modifying Protein 2 (RAMP2) and Calcitonin Gene-Related Peptide Type 1 Receptor (CALCRL), were quantified exclusively in SAdi. RAMP2 is a membrane-spanning protein that interacts with the receptor CALCRL enabling higher affinity binding of adrenomedullin (ADM) over calcitonin (McLatchie et al., 1998). We also observed that the orphan G protein-coupled receptor GPR182, which has been suggested although not confirmed to be part of the adrenomedullin signaling apparatus (Kapas et al., 1995; Kennedy et al., 1998), was only present in SAdi. These findings support previous evidence that Adrenomedullin regulates adipocyte beiging (Lv et al., 2016; Zhang et al., 2016). Consistent with the unique role of SAdi in beiging, we observed expression of a number of beiging-related proteins, such as CREB Regulated Transcription Coactivator 3 (CRTC3) (Shan et al., 2016; Xu et al., 2019), exclusively in SAdi. UCP1, a classic marker for adipocyte beiging, trended to be enriched in SAdi versus VAdi, although not significantly (**Fig. 2E**), likely because the mice from which these adipocytes were obtained were housed at room temperature. Strikingly, markers of adipose tissue myelinated neurons Myelin Protein Zero (MPZ) and Myelin Basic Protein (MBP) (Willows et al., 2021) were 320- and 21-fold higher in the SAdi proteome, respectively, highlighting greater neural innervation to SAdi (**Fig. 2D**), which is a major driver of adipocyte beiging (Blaszkiewicz et al., 2019).

We identified 69 proteins exclusively expressed in VAdi, and an additional 51 proteins enriched in adipocytes from this depot. This included the two ‘atypical’ subunits of extracellular matrix (ECM) protein collagen VI, Collagen alpha-5(VI) chain (COL6a5) and alpha-6(VI) chain (COL6a6), which were 28- and 5-fold higher in VAdi compared to SAdi, respectively (**Fig. 2F**). Interestingly, the typical collagen VI isoforms (COL6a1, COL6a2 and COL6a3) were not different between SAdi and VAdi (**Fig. 2F**). This suggests that there is a unique ECM composition between these different adipocytes even under chow-fed conditions, which involves the atypical, but not the typical, collagen VI isoforms. Also noteworthy, Cyclin-Dependent Kinase 6 (CDK6) was 15-fold more abundant in VAdi (**Fig. 2G**). CDK6 has been associated with expansion of adipocyte size (Sarruf et al., 2005) and suppression of adipose beiging (Hou et al. 2018) in part via crosstalk with the nutrient sensor mammalian Target of Rapamycin (mTOR). In line with this, Sodium-Coupled Neutral Amino Acid Transporter 9 (SLC38A9), which is a lysosomal amino acid transporter that communicates amino acid availability to mTOR to induce its activity (Rebsamen et al., 2015; Wang et al., 2015), was quantified exclusively in VAdi. These data indicate that these processes may operate synergistically to promote hypertrophy of VAdi during tissue expansion. To further investigate adipocyte size between the two different depots, we performed histological examination of fixed visceral and subcutaneous adipose depots in C57Bl/6J mice fed chow or WD for 8 weeks (**Fig. 3A-B**). This revealed that VAdi were significantly larger than SAdi in chow-fed mice and, upon WD feeding, underwent a 2.6-fold expansion in area compared to only a 1.7-fold expansion in SAdi, which supports the hypothesis that VAdi are primed to expand via hypertrophy. We next performed targeted analysis of exclusively or differentially expressed proteins between SAdi and VAdi from chow-fed mice in several metabolic pathways integral to adipocyte metabolism. Analysis of the glycolysis pathway revealed that the enolase class of enzymes were enriched in SAdi. ENO1 was exclusive to SAdi, ENO3 was significantly more abundant, and ENO2 trended higher (p = 0.052) in SAdi compared to VAdi. The enolase enzymes catalyze the interconversion of 2-Phosphoglycerate and Phosphoenolpyruvate in the penultimate step of glycolysis. As there were no other differences in levels of glycolytic proteins, including the insulin-responsive glucose transporter SLC2A4 (GLUT4), this may denote a separate function for enolase in adipocytes, as certain moonlighting functions for enolase have been described including stress response and tissue remodeling (Paludo et al., 2015). In contrast to differences in glucose metabolism, we observed no significant differences in the levels of proteins involved in lipid synthesis or storage (*i.e.* CD36, DGAT 1 and 2, LPL, FABP4, LDLR, PLN 1, 2 and 3, ACC1, ACLY, FASN and SCD2). Finally, both adipocyte types had similar levels of adipokines including Adiponectin (ADIPOQ), Leptin (LEP) and Resistin (RETN). Taken together, this comparison of SAdi and VAdi from lean mice reveals that these adipocyte proteomes are highly conserved, with depot-specific differences including select metabolic processes, extracellular matrix and regulators of cell size.

**Figure 3.**
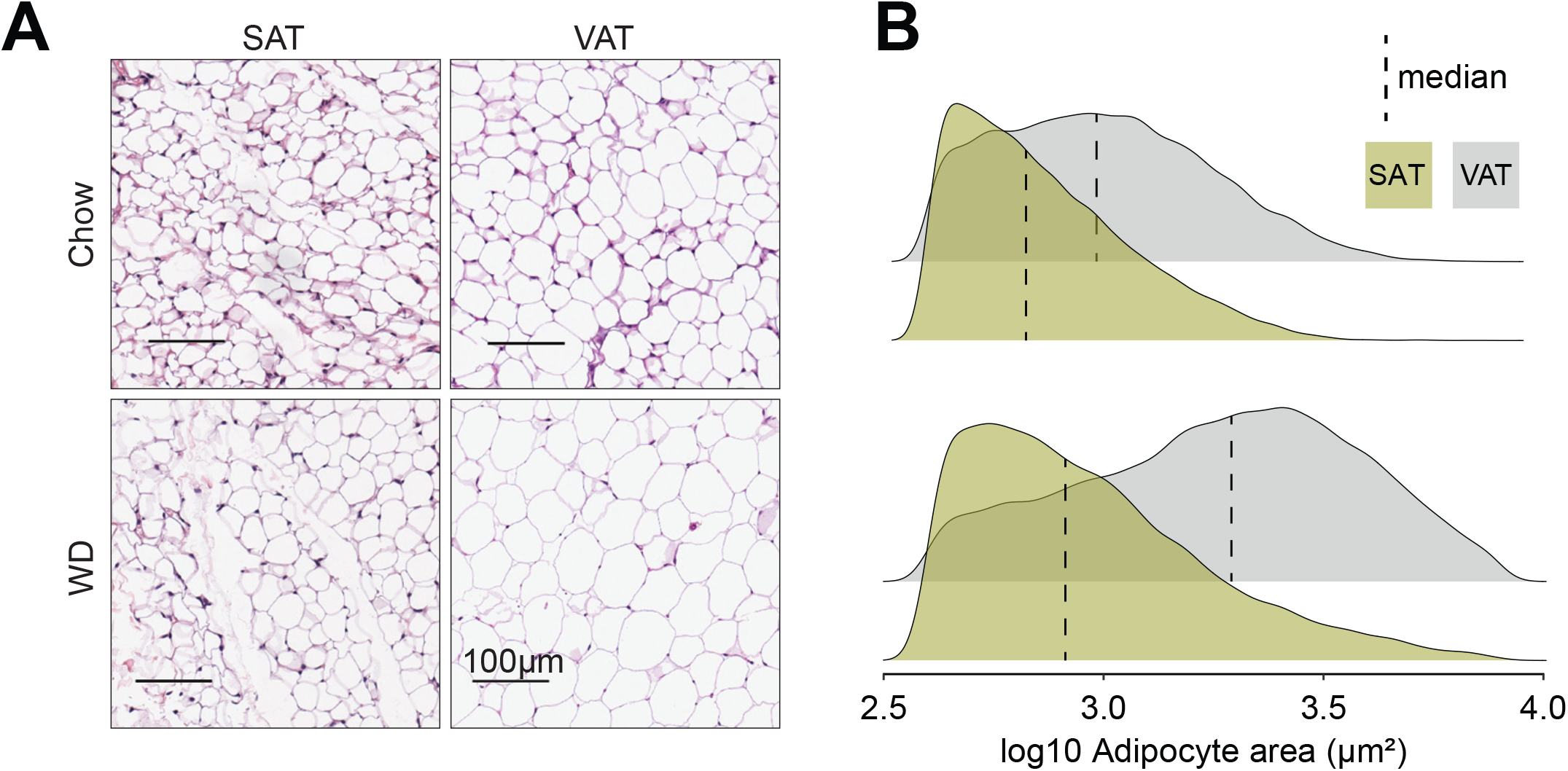
Visceral adipocytes are larger than subcutaneous and show greater expansion with WD feeding. **(A)** Representative H&E stained sections of fixed visceral and subcutaneous adipose tissue of male C57Bl/6J mice fed chow or WD for 8 weeks (scale bars represent 100 μm). **(B)** Density plots with medians of adipocyte areas pooled by depot and diet groups.

### VAT adipocytes are more diet responsive compared to SAT adipocytes

We next focused on diet-induced changes by comparing chow versus WD within each adipocyte type. Upon WD feeding, 16% (1046 proteins) of the total proteome changed in VAdi, in contrast to 7% (411 proteins) in SAdi (**Fig 4A-B**). There was striking overlap between the diet-regulated proteins in adipocytes between the different depots (54% of proteins) (**Fig. 4C**); thus, this conserved regulation likely defines core adaptive mechanisms across white adipocytes. Upregulated core proteins were enriched for ECM remodeling and organization (**Fig. 4D**), reflecting adipocyte hypertrophy. The majority of core regulated ECM remodeling proteins were also regulated in a depot-dependent manner (marked *) with greater expression in VAdi. Downregulated core proteins were enriched for fatty acid metabolism, cellular detoxification and branch chain amino acid catabolism. Coenzyme Q (CoQ) is a mitochondrial cofactor synthesized in mitochondria that is essential for mitochondrial respiration (Stefely and Pagliarini, 2017) and depletion of the CoQ biosynthesis machinery has been shown in a range of insulin resistance models in adipocytes (Fazakerley et al., 2018) and, notably, many proteins involved in mitochondrial CoQ biosynthesis (COQ4-7, 9, 10A and 10B) were decreased in adipocytes from both depots (**Fig. 4D**). Given the prominent role of mitochondria in insulin resistance (Diaz-Vegas et al., 2020) and the pronounced downregulation of the mitochondrial CoQ biosynthesis pathway (**Fig. 4D**), we next determine whether diet induced a selective or generalized depletion of mitochondrial proteins. Our adipocyte proteome covered 70% of the mitochondrial proteins described in the MitoCarta3.0 database (Rath et al., 2021). Sustained WD induced an overall decrease in mitochondrial protein abundance regardless of depot, with the majority of Complex I (NDUFA1-3;5-13,NDUFB2-11, NDUFS1-8, NDUFV1-3), Complex II (SDHA-D) and Complex III (Uqcrc1-2;11;B;C:FS1;H) downregulated and, to a lesser extent, Complex IV (COX4I1;6A1;6B1) and Complex V (ATPC1;5d;5O). Interestingly, a significantly larger proportion of mitochondrial proteins were decreased in VAdi compared to SAdi (31% vs. 15%, χ2 test, p<0.0001) suggesting that the mitochondrial proteome exhibits differential sensitivity to WD depending on depot (**Fig. 4E**). The ETC complexes reflected this relationship, as there was a striking reduction in proteins related to Complex I, II, III, and IV in VAdi after WD relative to SAdi on the same diet (**Fig. 4F**), suggesting that depletion of ETC subunits may contribute to mitochondrial dysfunction in VAdi. Since adipocyte death may be caused by excessive stress during obesity-related adipose tissue remodeling (Lee et al., 2010) we next explored the abundance of pro- and anti-apoptotic proteins in this dataset. Two stress-associated pro-apoptotic proteins which are localized to the outer mitochondrial membrane, BCL-2-Associated X (BAX) and BCL-2-Related Ovarian Killer Protein (BOK), were regulated in a depot-dependent manner in response to WD. We detected no change in expression of the counter-regulatory anti-apoptotic proteins (B-Cell Lymphoma 2 (BCL2) or MCL1 Apoptosis Regulator (MCL1)), indicating the vulnerability of VAdi from WD-fed mice to apoptotic signaling (**Fig 4G**). Interestingly, BAX and BOK have a role in mitophagy suggesting a possible interaction between mitophagy and cell death in adipocytes (He et al., 2021). Expression of the key adipogenic regulator Peroxisome Proliferator-Activated Receptor Gamma (PPARG) is decreased in hypertrophic adipocytes, which can cause mitochondrial dysfunction (Bond et al., 2019). We found that PPARG was regulated in a depot-dependent manner in response to diet, where PPARG was below the limit of detection in VAdi upon WD only, which points towards loss of adipocyte identity in VAdi (**Fig. 4H**). Furthermore, two targets of PPARG, BCL2/Adenovirus E1B 19 kDa Protein-Interacting Protein 3 (BNIP3) and the E3 ubiquitin ligase Membrane Associated Ring-CH-Type Finger 5 (MARCHF5), were regulated in a similar manner (**Fig. 4H**). Since BNIP3 is required for optimal mitophagy (Bellot et al., 2009), it is possible that BNIP3 downregulation induces the accumulation of dysfunctional mitochondria (Bond et al., 2019; Tol et al., 2016), increasing the susceptibility of VAdi to pro-apoptotic stimuli.

**Figure 4.**
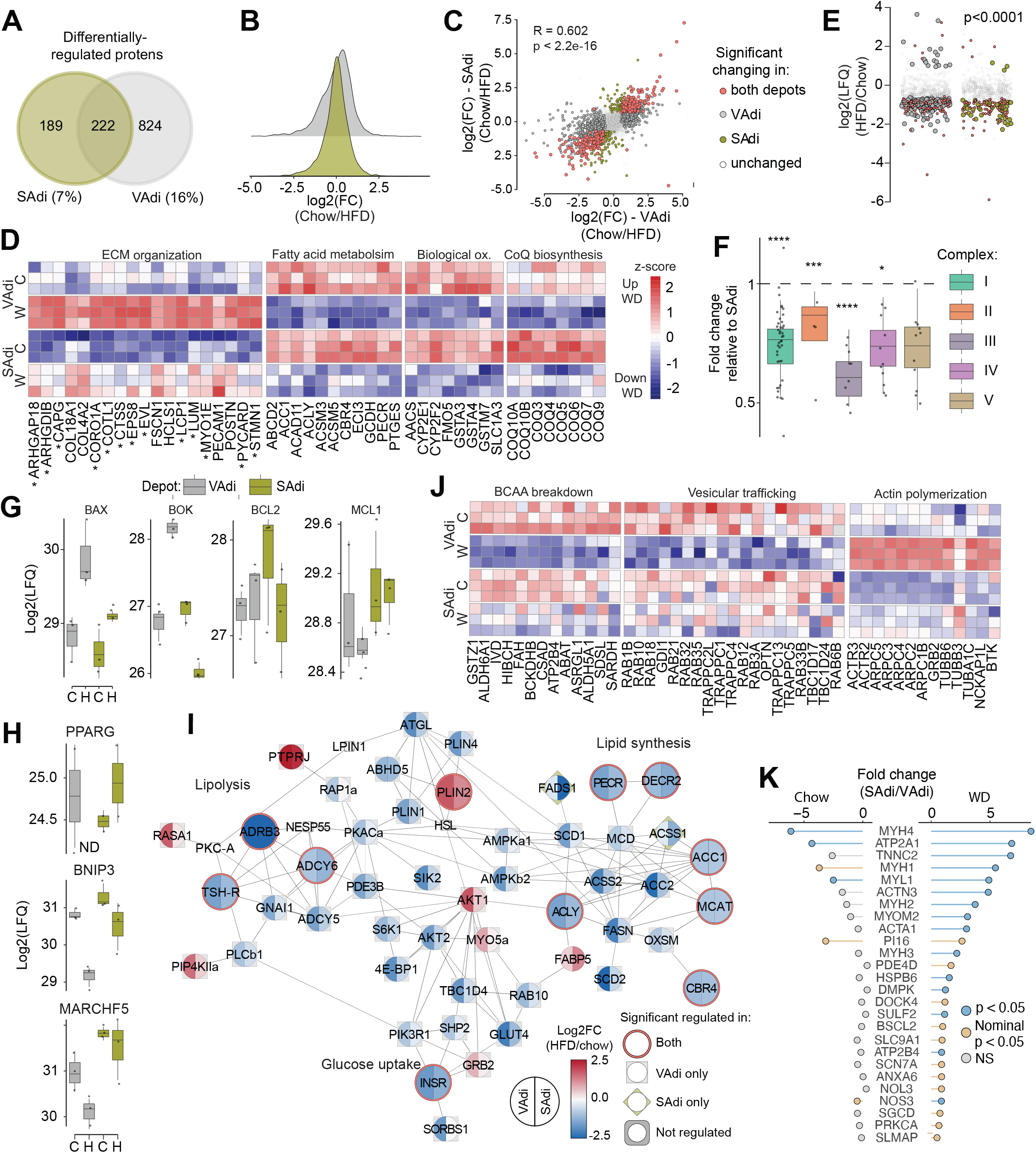
Adipocytes from the visceral depot are more diet-responsive than adipocytes from the subcutaneous depot. **(A)** Venn diagram showing differentially-regulated proteins in response to WD in isolated adipocytes from the two depots. **(B)** Distribution of log2 fold changes in response to WD in visceral and subcutaneous adipocytes (VAdi and SAdi, respectively). **(C)** Proteins abundances after WD across all proteins in VAdi (x-axis) and SAdi (y-axis) with correlation coefficient for direction and magnitude of changes. **(D)** Z-scored protein abundances of selected proteins within pathways that change in both depots in response to WD. **(E)** Changes in all mitochondrial proteins detected in VAdi or SAdi after WD. **(F)** Abundances of proteins in Complexes I-VI of the electron transport chain in VAdi after WD relative to SAdi. **(G-H)** Label-free quantitation for BCL-2-Associated X (BAX), BCL-2-Related Ovarian Killer Protein (BOK), Apoptosis regulator Bcl-2 (BCL2), Myeloid leukemia cell differentiation protein Mcl-1 homolog (MCL1), Peroxisome proliferator-activated receptor gamma (PPARG), BCL2/adenovirus E1B 19 kDa protein-interacting protein 3 (BNIP3) and E3 ubiquitin-protein ligase MARCHF5 (MARCHF5). **(I)** Z-scored protein abundances of selected proteins within pathways uniquely changing in VAdi in response to WD. **(J)** Protein differences in smooth-muscle proteins in VAdi and SAdi in mice fed either chow or WD. C: Chow, W: Western diet

PPARG is a major regulator of lipid metabolism and glucose homeostasis; therefore, we further interrogated these processes by constructing a protein-protein interaction network of proteins involved in lipolysis, lipid synthesis and storage, and glucose uptake, then filtered these for diet-responsive proteins in VAdi or SAdi (**Fig. 4I**). Key receptors modulating glucose and fat metabolism, including the Insulin Receptor (INSR) and the Beta-3 Adrenergic Receptor (ADRB3), were downregulated in adipocytes from both depots after sustained WD feeding. However, several additional key metabolic proteins were downregulated selectively in VAdi, including the lipolytic proteins Phosphodiesterase 3B (PDE3B; 3-fold) and Adipose Triglyceride Lipase (ATGL; 4-fold), proteins involved in the glucose transport pathways such as GLUT4 (4-fold), RAB10 (2-fold) and TBC1D4 (3-fold). The lipogenic proteins Fatty Acid Synthase (FASN; 4-fold) and Acetyl-CoA Carboxylase 2 (ACC2; 5-fold) were also downregulated in VAdi, whereas we did not observe diet regulation of fatty acid transport proteins including CD36, FABP4 or LDLR. Further pathway analysis of proteins that were regulated by WD specifically in VAdi revealed a decrease in proteins involved in branched-chain amino acid catabolism and several Rab GTPase proteins and other vesicle trafficking proteins and increased actin polymerization (**Fig. 4J**). The VAdi from WD mice was also enriched in proteins involved in inflammation and immune-related processes, such as ‘leukocyte proliferation’ and ‘adaptive immune response’. This was surprising, as the adipocytes were isolated by collagenase digestion and flotation. One possibility is that fat-laden immune cells, such as macrophage foam cells, co-fractionate with adipocytes during flotation. It should be noted that similar enrichment of immune-related proteins in VAdi has been reported by others (Jones et al., 2020). Our observations in VAdi of increased expression of cytoskeletal and ECM proteins and decreased expression of proteins specific to adipocyte function, together with a decrease in PPARG, reveal a transition of VAdi to a fibroblast-like state.

Interestingly, we identified several proteins enriched in the SAdi proteome from WD-fed animals which are typically associated with smooth muscle function, such as Myosin Light Chain 1 (MYL1), Myosin Light Chain 4 (MYL4), Myosin-4 (MYH8), Skeletal Muscle Actin Alpha 1 (ACTA1), Alpha-Actinin-3 (ACTN3), Troponin T2 (TNNT2) and the sarcoplasmic/endoplasmic reticulum calcium ATPase 1 (ATP2a1), with differences in expression ranging from 6 to 120-fold compared to VAdi from the same animals. This was specifically an adaptation to WD, as this disparity was not observed between adipocytes from chow-fed animals (**Fig. 4K**). The enrichment of muscle-associated proteins in SAT is consistent with this depot’s unique Prx1-expressing smooth muscle lineage (Sanchez-Gurmaches et al., 2015), and this signature resembles that of beige adipocytes residing in SAT (Long et al., 2014). A mechanical actomyosin response is important for adrenergic stimulation of brown adipocytes to maintain their stiffness and uncoupling capacity (Tharp et al., 2018), suggesting this is an important mechanism for SAdi to maintain cytoskeletal stiffness independent of the ECM during nutrient overload. Recently, it was speculated that these proteins are involved in adipose tissue innervation, as overexpression of the fat-derived neurotrophic factor Neurotrophin 3 increases adipose tissue innervation and a subsequent increase in striated muscle specific proteins (Cui et al., 2021), so this may reflect differences in neural innervation between depots to confer beta-adrenergic responsiveness.

### Whole-tissue proteomics in VAT, but not SAT, reflects changes to the adipocyte proteome upon WD feeding

Adipose tissue contains numerous cell types in addition to adipocytes (Emont et al., 2022). Cell type deconvolution of subcutaneous and visceral depots at the whole-tissue level (SAT and VAT, respectively) utilizing an adipose single nuclei transcriptome database (Emont et al., 2022; Jew et al., 2020) (**Fig. 5A**) revealed that preadipocytes, macrophages and adipocytes were the predominant cell types, with the only difference between the depots being significantly higher dendritic cells in VAT. WD was associated with a significant increase in macrophages and a decrease in adipocytes in VAT, but not SAT (**Fig. 5A**). To more closely assess diet-induced changes in the two tissues, we selected tissue-enriched proteins (>1.5 fold higher expression in tissue, p < 0.01) or proteins exclusively expressed in tissue versus adipocytes. Of these, we identified 56 tissue-enriched proteins in SAT and 91 in VAT that had increased expression with WD. Seven proteins were upregulated in both tissues, representing conserved mechanisms for adipose tissue adaptation to WD. These included proteins involved in cytoskeletal organization (SSH3, CDAN1 and SCAMP2), ECM formation (PXDN1), estradiol metabolism (CYP1B1), lipoprotein metabolism (APOA4), and a DNA polymerase subunit (POLE3). In SAT, 84 proteins were upregulated specifically in response to WD, with particular enrichment of the E3 ubiquitin ligase pathway (RNF146, UBE2W, FBXO7 and FBXW17). In VAT, subunit 2 of the NFκB complex (NFKB2) was upregulated with WD, as well as BCL10, which is essential for lymphocyte-induced activation of NFkB. NFκB activation in adipose tissue is important for recruitment of immune cells (Griffin, 2022; Hill-McAlester, 2015), and may represent a depot-specific mechanism for the observed diet-induced recruitment of macrophages in VAT.

**Figure 5.**
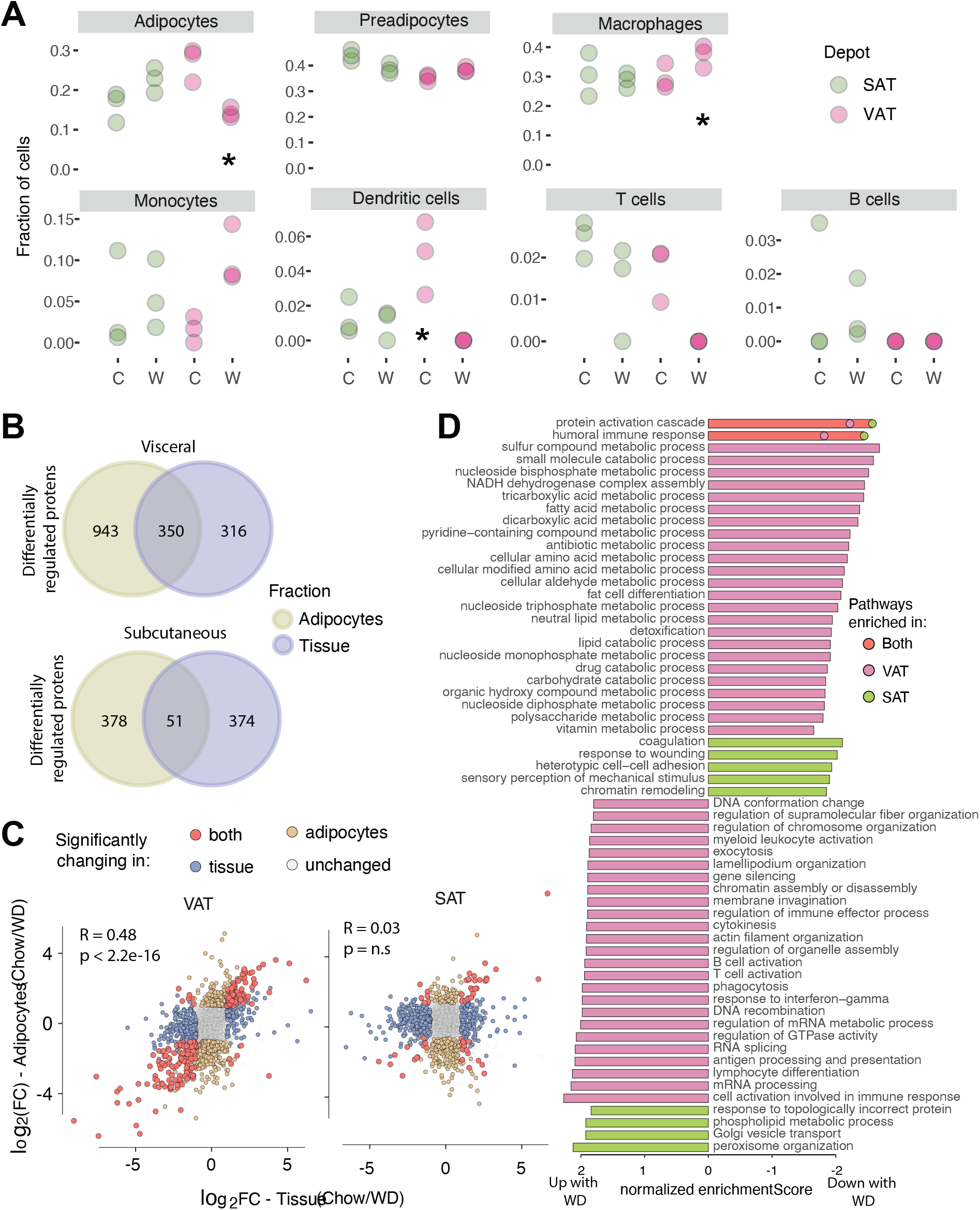
WD causes a shift in visceral adipose tissue toward macrophage infiltration and fibrosis, and WD adaptations in subcutaneous adipose tissue occur primarily in non-parenchymal cells. **(A)** Cell type deconvolution using proteomic data from subcutaneous or visceral whole tissue of mice fed chow (C) or Western diet (W). * indicates significant difference by 2-way ANOVA with Tukey’s post hoc test; p < 0.05. **(B)** Overlap of proteins changing with Western diet (WD) in isolated adipocytes and whole-tissue in the visceral or subcutaneous depots. **(C)** Correlation of changes in protein abundance after WD across all proteins in tissue and adipocytes within each depot. **(D)** Gene ontology of proteins changing with WD in visceral and subcutaneous adipose tissue (VAT and SAT, respectively) (showing pathways which are significant after false discovery correction p < 0.05).

Due to the difficulty of isolating primary adipocytes, many studies only examine changes in the whole tissue and extrapolate these to estimate changes in adipocytes. In examining the validity of this approach, we observed that the overlap between diet-responsive proteins from VAdi and VAT was 350 proteins (**Fig. 5B**). In the case of the visceral depot, the direction and magnitude of changes in response to diet between adipocytes and whole tissue were highly correlated (R = 0.48 p<2.2e-16) (**Fig. 5C**), indicating that VAdi adaptations to WD could be reasonably detected at the tissue level. Pathways that were commonly upregulated between VAdi and VAT included cell growth *(e.g.* pathways associated with mRNA processing and actin filament organization) and immune system (**Fig. 5D**). Commonly down-regulated pathways included fatty acid and lipid metabolic processes, fat cell differentiation, amino acid metabolic processes and energy-related pathways (**Fig. 5D**). In the case of the subcutaneous depot, however, only 51 diet-regulated proteins overlapped between SAdi and SAT (**Fig. 5B**) and there was no correlation in diet response (**Fig. 5C**). These data suggest that the overall proteomics changes at a tissue level within SAT does not reflect the changes to adipocytes of the same tissue.

### Proteome comparison of the 3T3-L1 adipocyte model with primary adipocytes

3T3-L1 adipocytes are the standard model system to study adipocyte biology. Based on the depth of our murine proteome (**Fig. 1B**) we reasoned it would be valuable to compare these with the 3T3-L1 adipocyte proteome to determine the robustness of this model. To this end, we generated a deep proteome containing 8,862 proteins across four biological replicates of 3T3-L1 adipocytes. 90% of proteins identified in the isolated adipocyte proteome were identified in the 3T3-L1 proteome. The intensity Based Absolute Quantification (iBAQ) values of the overlapping proteins were strongly correlated between adipocytes from both VAT and SAT (R = 0.75; p < 2.2e-16 for VAdi; R = 0.73; p < 2.2e-16 for SAdi), indicating similar relative expression levels on an individual protein basis. Thus, many important adipocyte-specific processes were highly conserved in the 3T3-L1 adipocyte model at the proteome level.

Given the high concordance between 3T3-L1 and isolated adipocyte proteomes, we next ascertained differences that may be important to consider when translating findings from 3T3-L1 adipocytes to primary adipocytes. We designated proteins with less than 5-fold difference in expression in primary adipocytes versus 3T3-L1 adipocytes to be ‘normal range’, as the abundances of 99.5% of the proteins in the VAdi and SAdi proteomes from chow-fed mice were within 5-fold (**Fig. 2B**). Of proteins identified in both the 3T3-L1 and primary adipocyte proteomes, 75% of proteins were within a 5-fold range (**Fig. 6A-B**), indicating good conservartion between the two proteomes. However, pathway analysis revealed that there was a clear enrichment of ETC proteins and especially for proteins within Complex I in isolated adipocytes independent of depot (**Fig. 6C**). Furthermore, a number of immune related processes were enriched in the isolated adipocyte together with proteins promoting angiogenesis, Annexin A1 and A3, fibroblast growth factor 1 and 2 and integrin beta 1. We also observed several pathways intrinsic to adipocyte biology that were not conserved in 3T3-L1 adipocytes, such as fatty acid and glycerolipid metabolism, including FABP4, Phosphoenolpyruvate Carboxykinase 1 (PEPCK1), CD36, PLIN1, and ATGL. Notably, Leptin was exclusively expressed in primary adipocytes, and Resistin was 20 to 30 times more abundant compared to 3T3-L1 adipocytes. Thus, the increased expression of these proteins in primary adipocytes correlates with the higher lipid content and larger, unilocular lipid droplets compared to 3T3-L1 adipocytes (Olson, 2018). Lastly, a number of proteins within cAMP signaling, most notably exemplified with the beta 3 adrenergic receptor, were highly enriched in isolated adipocytes. In contrast, proteins with higher abundance in 3T3-L1 adipocytes were enriched for chromatin assembly, ribosome assembly and RNA splicing pathways, features which are characteristic of proliferative cells. These data highlight potentially important considerations when using the 3T3-L1 adipocyte model. However, overall, the 3T3-L1 adipocyte proteome is highly representative of primary adipocytes.

**Figure 6.**
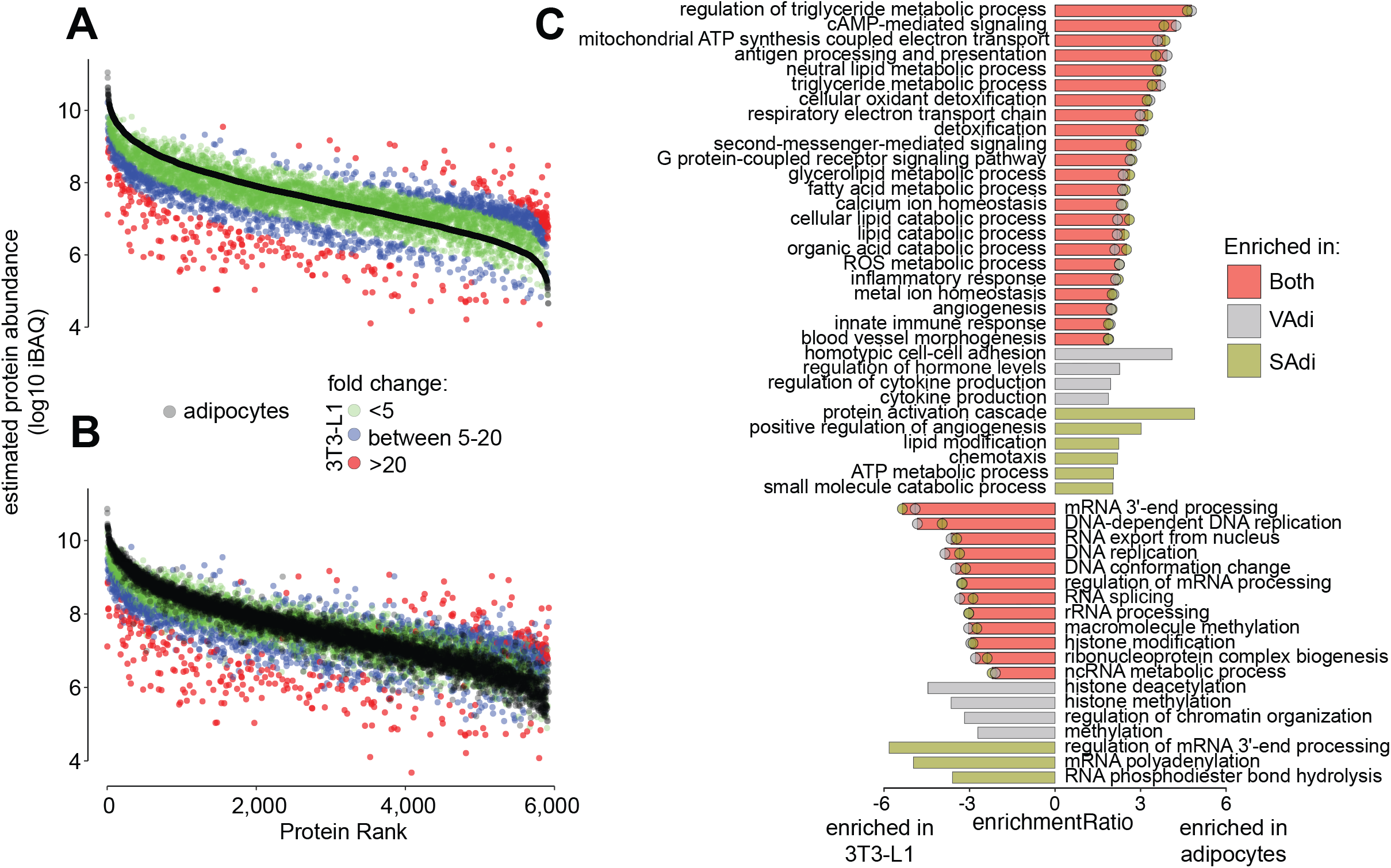
Comparison of the 3T3-L1 and primary murine adipocyte proteomes. **(A-B)** Ranked protein intensities based absolute quantification (iBAQ) of 3T3-L1 adipocytes and **(A)** visceral adipocytes (VAdi) or **(B)** subcutaneous adipocytes (SAdi) from chow-fed C57Bl/6J mice. Colors denote fold changes in protein abundance between primary murine adipocytes and 3T3-L1 adipocytes. **(C)** Gene ontology of proteins that were differentially-regulated between 3T3-L1 and primary murine adipocytes.

## DISCUSSION

Subcutaneous and visceral adipose tissues have well-documented differences that have been implicated in their differential association with metabolic disease. This study presents a global and unbiased proteomic analysis of these two different depots in order to define the molecular features that might govern these functional differences. These data should serve as an invaluable resource for researchers interested in adipose biology.

By studying the proteomes of isolated adipocytes from visceral and subcutaneous depots, we have found that the major depot-specific differences are encoded by just 3% of the proteome. Nevertheless, several major biological processes are encoded within this small subdomain of the proteome. These include various components of the mTORC1 pathway such as SLC38A9 and CDK6, and cytoskeletal components such as COL6a5 and COL6a6 that are enriched specifically in VAdi. COL6a5 and COL6a6 atypical collagens can replace the typical collagen VI isoforms (a1, a2 and a3) in the collagen VI monomers. Collagen VI is implicated in a deleterious fibrotic response in adipose (Beals et al., 2021; Khan et al., 2009), and inhibition of COL6a5 is associated with improvements in lipid metabolism in adipocytes (Sun et al., 2021). We observe that visceral adipocytes in C57Bl/6 mice are significantly larger than those from the subcutaneous depot, and mTOR is known to play a role in controlling adipocyte cell size (Chakrabarti et al., 2010; Lee et al., 2016). Furthermore, larger cells require profound shifts in cytoskeleton to support size expansion. Conversely, SAdi were enriched in mitochondrial proteins, particular proteins that comprise Complex I of the ETC, indicating gearing of energy machinery towards carbohydrates as a fuel source (Kappler et al., 2019) and possibly rendering SAdi more metabolically flexible (Goodpaster and Sparks, 2017). Taken together with the trend for increased expression of major lipolysis and beiging proteins in SAdi, these processes are likely coordinately regulated and complementary to subcutaneous adipocyte metabolism.

In contrast to visceral, subcutaneous adipocytes are capable of undergoing browning or beiging. One of the signature features of this process is the increased expression of the uncoupling protein UCP1, which provides the molecular basis for increased thermogenesis. Interestingly, we observed several other molecular features that may participate in this unique beiging process in subcutaneous adipocytes. For example, an enrichment in muscle-associated actomyosin proteins, which are important for the mechanical aspect of adrenergic stimulation (Tharp et al., 2018) which controls lipolysis and beiging; increased expression of proteins involved in neural innervation exemplified with structural components of myelin sheath of neurons (MPZ and MBZ) (Nguyen et al., 2014; Willows et al., 2021), and enrichment of several subunits of the adrenomedullin receptor including RAMP2 and CALCRL (Lv et al., 2016). Importantly, RAMP2 expression is associated with beneficial metabolic effects, as single nucleotide polymorphisms in the RAMP2 gene have been associated with changes in BMI and T2D in humans, and ADM has been shown to improve insulin sensitivity when administered to WD-fed mice (Zhang et al., 2016). Taken together, we observed few, but potentially important, differences that define lean adipocytes between these two depots.

Both adipose depots underwent profound changes in a sustained obesogenic environment by WD feeding. We combined our whole-tissue proteomics data with single-RNA sequencing data from murine adipose tissue (Emont et al., 2022; Jew et al., 2020) to achieve a more holistic overview of these depots. This approach shows that VAT increased in both preadipocytes and immune cells (macrophages) in response to sustained WD feeding. These data are in line with the ‘adipose tissue expandability model’, which identifies limited lipid storage capacity during energy surplus followed by adipose tissue dysfunction (Pellegrinelli et al., 2016), which has been reported previously in C57Bl/6 mice (van Beek et al., 2015). Furthermore, by overlaying the proteomics changes with diet in whole adipose tissue and isolated adipocytes, we uncovered concordance between adipocytes and tissue only in the visceral depot. This is an important observation for two reasons. First, this distinction is crucial for future studies of adipose tissue biology, as assumptions of adipocyte biology may be distorted by whole tissue approaches, and second, dietary responses in subcutaneous tissue cannot be explained by the adipocytes themselves, warranting future research. Notably, a large part of the proteomic adaptations to diet within SAdi also occur in VAdi, which we define as a set of core additive proteins albeit VAdi displayed a much greater diet response than SAdi. Mitochondrial dysfunction is thought to be a hallmark of adipose dysfunction (Diaz-Vegas et al., 2020; Kusminski and Scherer, 2012). During mitochondrial stress, both apoptotic signaling and mitophagy is activated, where enhanced mitophagy facilitates cell survival by removing damaged mitochondria(Kubli and Gustafsson, 2012). Intriguingly, a large part of the mitochondrial proteome was decreased with sustained WD in adipocytes independent of depot. However, this decrease was greater in VAdi with a concurrent increase in apoptotic signaling and a decrease in the level of a key regulator of mitophagy, BNIP3 (Bellot et al., 2009), pointing towards severe mitochondrial stress in VAdi. Interestingly, PPARG was selectively downregulated in VAdi upon WD together with many proteins involved in highly adipocentric pathways, such as lipid breakdown and storage and glucose uptake, whereby VAdi take on a fibroblast-like cell identity. PPARG regulates BNIP3 (Tol et al., 2016) and, in line with this, PPARG agonists ameliorate mitochondrial function in obese white adipose tissue (Choo et al., 2006). These data highlight a depot-dependent protein fingerprint in response to WD, where VAdi showed signs of mitochondrial dysfunction, possibly through downregulation of PPARG.

This analysis also provided an opportunity to investigate the suitability of 3T3-L1 adipocytes as a model of adipocyte biology by proteomics profiling. Strikingly, 3T3-L1 and isolated adipocytes share ~90% overlap at the proteome level supporting the robustness of this in vitro cell mode and their proteomes were highly correlated. However, there were some key differences; first, the isolated adipocytes were enriched in proteins related to organismal biology, highlighted by immune related proteins, but also proteins involved in angiogenesis stating that isolated adipocytes receive cues from multiple cell types. Second, isolated adipocytes were enriched in adipocyte proteins, such as ATGL, PLIN1 and the beta 3 adrenergic receptor, and third, 3T3-L1 protein profile point towards a more proliferative cell type. The last two presumably work cooperatively giving rise to larger lipid droplets in isolated adipocytes.

This comparison of the adipocyte-specific and adipose tissue proteomes from lean and obese mice has revealed that the adipocytes from subcutaneous and visceral white adipose depots are profoundly similar. However, we identified several important biological processes unique to each adipocyte type, which are likely important regulators of adaptation to WD and resultant systemic effects associated with the different adipose depots. Notably, we observed unique tissue-specific processes which support the functions of the resident adipocytes and likely drive many unique adipose depot functions. This deep proteomics analysis will serve as a resource to complement other studies that have compared the proteomes or transcriptomes of different adipose tissue depots from mice or humans to better understand adipose biology.

## Author Contributions

DEJ, SJH and VD conceived the study. VD, KCC, AH, JS performed animal experiments. SJH and VD performed LC-MS experiments. SM, MEN, ADV and JGB anlysed the data. SM, MEN and DEJ wrote the manuscript. All authors agreed on the final version of the manuscript.

## Acknowledgments

We thank SydneyMS for providing the instrumentation used in this study and the Laboratory Animal Services at the Charles Perkins Centre of the University of Sydney. EchoMRI was performed in the Sydney Imaging Facility, University of Sydney. Histology was performed in the Histopathology Facility of the Garvan Institute of Medical Research. This work was supported by Australian Research Council project grant FL200100096 (to DEJ).

---

**Supplemental Figure 1.**
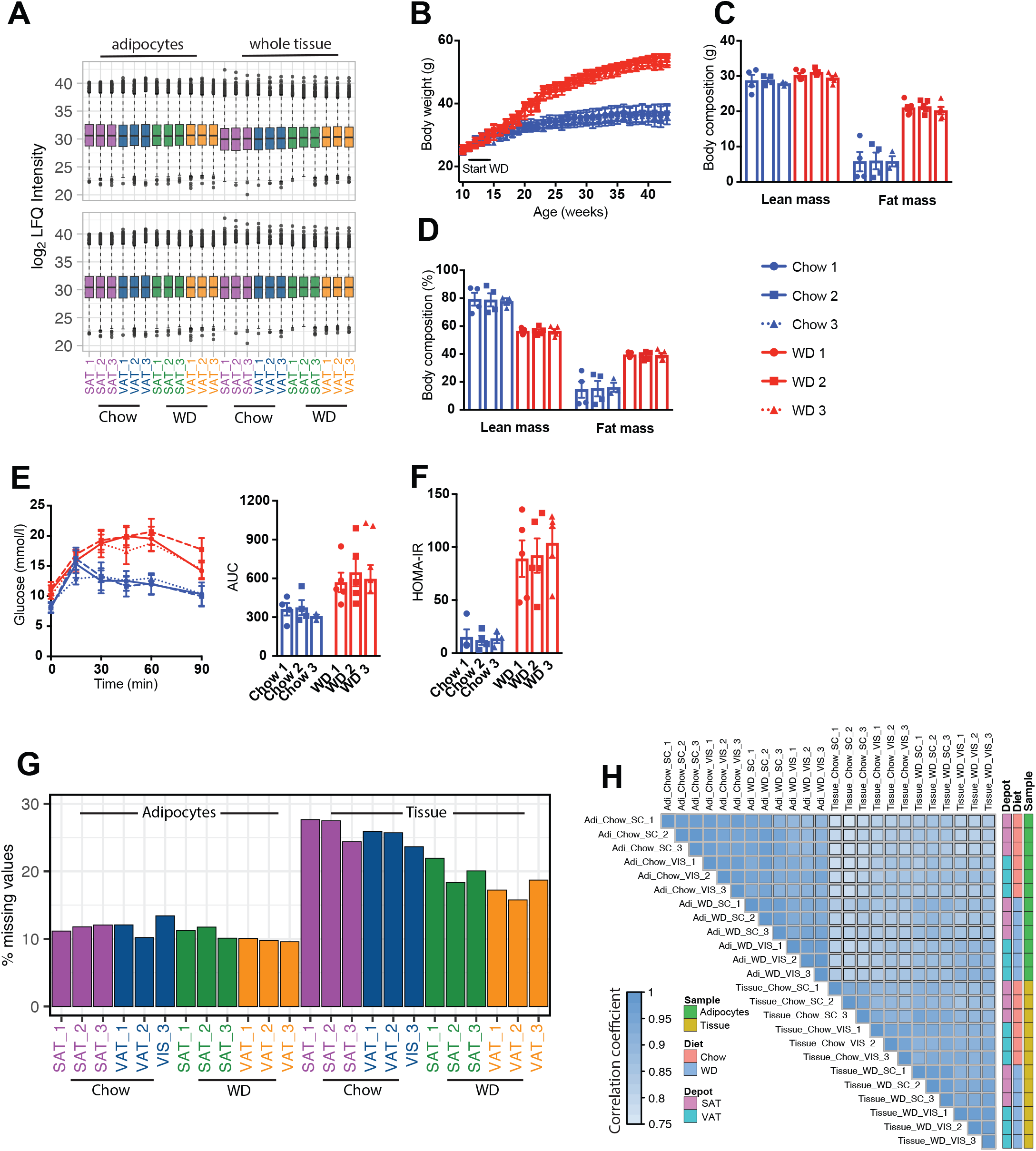
**(A)** Box plot of all proteomics samples before (top) and after (bottom) median normalization. **(B)** Body weights of mice fed chow of WD for 9 month. **(CD)** Body composition in grams **(C)** and percent **(D)** at 9 month of diet. **(E)** Blood glucose and area under the curve during a glucose tolerance test. **(F)** Homeostatic Model Assessment for Insulin Resistance (HOMA-IR) from fasting blood glucose and plasma insulin levels. **(G)** Percent missing in proteomics data across all samples. **(H)** Correlation coefficient for proteomics across all samples.

